# Estimating Rates of Progression and Predicting Future Visual Fields in Glaucoma Using a Deep Variational Autoencoder

**DOI:** 10.1101/652487

**Authors:** Samuel I. Berchuck, Sayan Mukherjee, Felipe A. Medeiros

## Abstract

**Purpose:** To develop a novel deep learning algorithm to improve estimation of rates of progression and prediction of future patterns of visual field loss in glaucoma.

**Design:** Prospective observational cohort.

**Methods:** A variational auto-encoder (VAE) was trained to learn a low-dimensional feature representation of standard automated perimetry (SAP) visual fields using 29,161 fields from 3,832 patients. The VAE was trained on a 90% sample of the data, with randomization at the patient level. Using the remaining 10%, rates of progression and predictions were generated, with comparisons to SAP mean deviation (MD) rates and point-wise (PW) regression predictions, respectively. From the VAE, rates were calculated using the average of slopes across latent features from ordinary least squares (OLS) regression and trajectories of the features were used to generate predictions.

**Results:** The longitudinal rate of change through the VAE latent space (e.g., with eight dimensions) detected a significantly higher proportion of progression than MD at two (19% vs. 6%) and four (40% vs 14%) years from baseline. Early on, VAE improved prediction over PW, with significantly smaller mean absolute error in predicting the 4^th^, 6^th^ and 8^th^ visits from the first three (e.g., visit eight: VAE8: 4.06 dB vs. PW: 6.06 dB; P<0.001).

**Conclusion:** A deep VAE can be used for assessing both rates and trajectories of progression in glaucoma, with the additional benefit of being a generative technique capable of predicting future patterns of visual field damage in the disease.

## INTRODUCTION

Glaucoma is a progressive optic neuropathy that results in characteristic changes to the optic disc and retinal nerve fiber layer.^1^ Although damage from glaucoma is irreversible, early treatment can usually prevent or slow down progression to functional damage and visual impairment.^2^ Assessment of functional damage is essential for management of the disease and standard automated perimetry (SAP) remains the default method for monitoring functional changes in the disease.^3^

Estimation of rates of functional deterioration by SAP is essential for determining patient prognosis and aggressiveness of therapy. However, the best method for estimating such rates is still a matter of controversy.^4^ While rates of change can be estimated by global parameters, such as mean deviation (MD), these rates would potentially ignore fast localized losses occurring in an otherwise mostly stable field. Several previous studies have approached this problem by attempting to project the high-dimensional (i.e., multiple locations) visual field to a lower dimensional representation, thus reducing variability and removing collinearity in the visual field measurements. This has been implemented successfully using a variety of unsupervised dimension reduction techniques^5–8^, that learn latent features derived from the original visual field data. Therefore, from the original 52-dimensional visual field on the SAP 24-2 test, for example, these techniques arrive at a much lower number of “meaningful” latent dimensions summarizing the original visual field.

Rates of progression can then be derived by analyzing the slopes of latent features across time. However, although previously used techniques may be able to learn a latent feature space; they cannot be used as realistic generators of visual field data, as they are designed explicitly for dimension reduction. Besides estimating how fast a patient is progressing (i.e., the current rate of change), it is also important to be able to generate, or predict, future visual field data from the currently available data for that patient. Such predictions would carry important prognostic value, by determining the likely areas of future damage over time which can help inform the impact of the disease on quality of life^9^.

In contrast to previously used models, a generative model learns a data generating mechanism (i.e., distribution), thus allowing for predictions of future visual fields. Historically, the task of learning a distribution of a high-dimensional data object was untenable without making simplifying assumptions, leading to the use of techniques like point-wise (PW) regression for visual fields.^10^ Modeling high-dimensional processes has become a reality with the explosion of deep learning. Deep learning has been applied effectively in health application, including the field of ophthalmology, ^11–13^ and in particular for visual fields^14, 15^. The most common generative models are the generative adversarial network and variational auto-encoder (VAE)^16, 17^. These methods can learn the complex distributions of high-dimensional biological data by first learning a projection to a lower-dimensional latent space, and then a mapping back to the original image. The latent space of these generative models has previously been shown to reveal novel biological patterns.^18, 19^ In this manuscript, we implement a VAE to learn a clinically motivated latent representation which can be used to determine rates of disease progression, and consequently, a generative process for visual field data.

## METHODS

Visual fields included in this analysis came from participants enrolled in a prospective longitudinal study designed to evaluate functional impairment in glaucoma. Written informed consent was obtained from all participants and the institutional review board and human subjects committee approved all methods. All methods adhered to the tenets of the Declaration of Helsinki for research involving human subjects and the study was conducted in accordance with the regulations of the Health Insurance Portability and Accountability Act.

During follow-up, patients underwent comprehensive ophthalmologic examinations, including review of medical history, visual acuity, slit-lamp biomicroscopy, intraocular pressure measurement, gonioscopy, dilated funduscopic examination, stereoscopic optic disc photography, and SAP using 24-2 Swedish interactive threshold algorithm standard (Carl Zeiss Meditec, Inc, Dublin, California, USA). Visual fields were excluded in the presence of eyelid or rim artifacts, fatigue effects, or evidence that the visual field results were caused by a disease other than glaucoma. Visual fields were also excluded if they had more than 33% fixation losses or more than 15% false-positive errors.

The VAE is an unsupervised technique, however in order to visualize the latent feature space of the VAE, we defined disease status as normal, suspect, and glaucoma. Glaucoma was defined by the presence of two or more repeatable glaucomatous visual field defects at baseline, defined as a pattern standard deviation with *P* < 0.05, or a Glaucoma Hemifield Test result outside normal limits, and corresponding optic nerve damage. Eyes were considered normal if they were recruited from the general population and had no visual field defects. The remaining eyes were considered suspects and had history of high intraocular pressure or suspicious glaucomatous appearance of the optic nerve, but in the absence of confirmed visual field defects.

### Visual Field Preparation

The 24-2 SAP produces visual fields with 52 informative data observations that together constitute an eye’s field of vision. The shape of a visual field is not rectangular, as is typically required for deep learning models, so square images were created by padding the visual field with zeros. As a result, the visual field images in our analysis were transformed to 12 × 12 squares (see Figure 1A for an example visual field used in the analysis). To prepare the visual fields for the deep learning algorithm, all visual field images were converted to right eyes for uniformity. In this study, we represented functional loss using total deviation (TD) values, an age-adjusted measure of sensitivity loss, measured in decibels (dB). TD is a continuous measure, with large negative values indicating functional loss.

**Figure 1.**
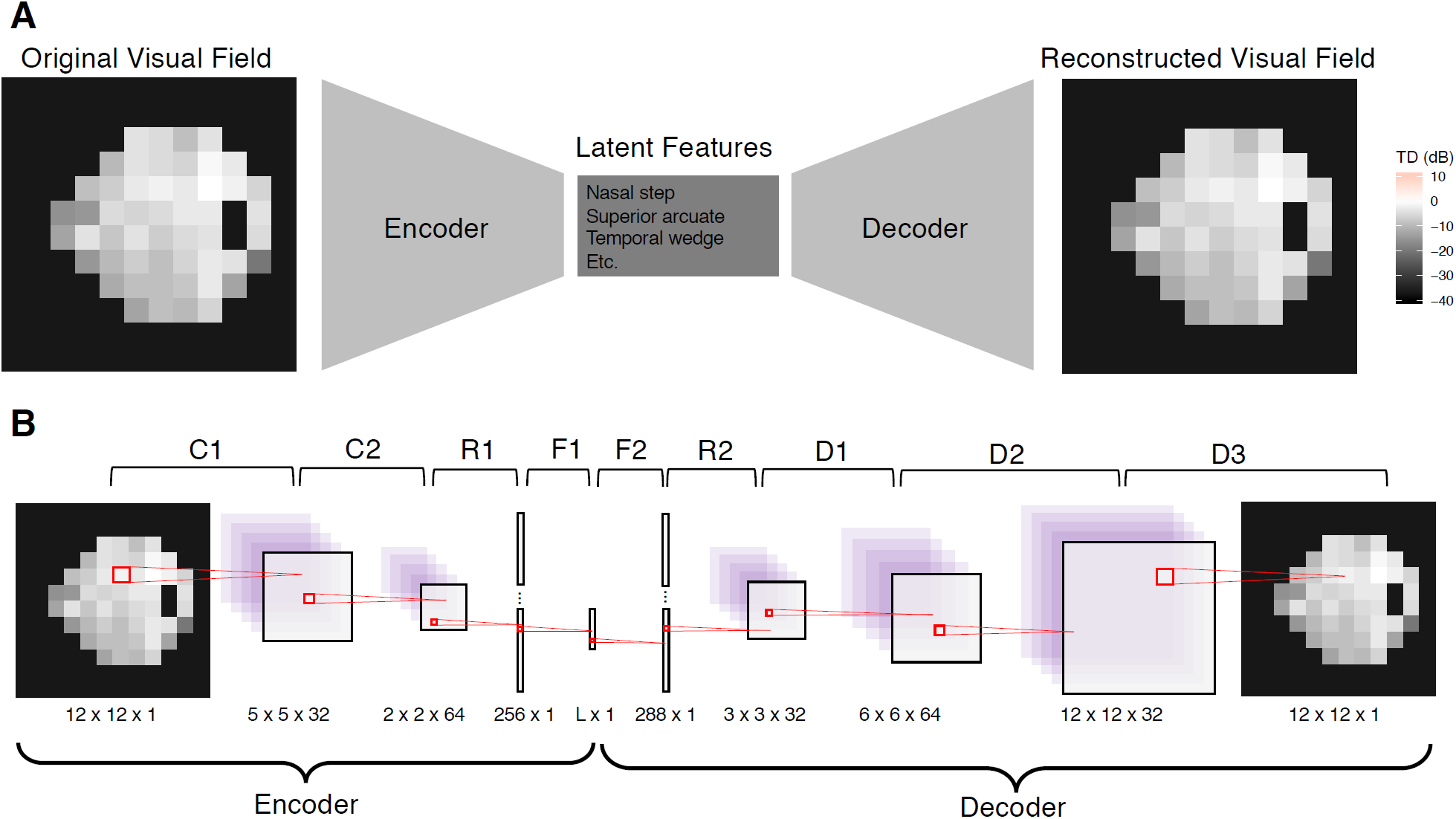
Illustrating the VAE for visual fields. A. The top frame demonstrates the structure of the VAE, which is a dual-mapping from the original visual field to the latent features and then back to a reconstructed visual field. B. The bottom frame provides details on the network architecture. There are four types of layers: convolutional (C), resizing (R), fully connected (F), and de-convolutional (D). The sizes at the bottom of each image reflect the transformed dimensions after each layer. For each layer, the kernel is 3×3 with a stride of 2. The activation function for all layers is the rectified non-linear unit, except for F1 and D3, which have the identity and sigmoid activations, respectively. The latent dimension, L, is a user specified parameter.

Prior to analysis, all visual fields were normalized to be in the range of zero and one. The training, validation, and test datasets were created by randomly sampling patients from the overall study population with 80%, 10%, and 10% probability, respectively. The randomization process was performed at the patient level, so that all images from a patient are included in at most one dataset. The demographics of the study datasets are illustrated in Table 1.

**Table 1.** Demographic and clinical characteristics of subjects included in the study across training, validation, and test data sets.

|  | **Total** | **Machine Learning Datasets** |  |  |
| --- | --- | --- | --- | --- |
|  |  | **Training** | **Validation** | **Test** |
| Number of Visual Fields | 29,161 | 23,744 (81%) | 2,577 (9%) | 2,840 (9%) |
| Number of Patients | 3,832 | 3,108 (81%) | 363 (9%) | 361 (9%) |
| Normal (n, %) | 667 (17%) | 547 (18%) | 58 (16%) | 62 (17%) |
| Suspect (n, %) | 2,221 (58%) | 1,793 (58%) | 222 (61%) | 206 (57%) |
| Glaucoma (n, %) | 944 (25%) | 768 (25%) | 83 (23%) | 93 (26%) |
| Number of visits | 7.61 (7.35) | 7.64 (7.36) | 7.10 (7.24) | 7.87 (7.29) |
| Follow-up period (years) | 4.95 (5.25) | 4.94 (5.24) | 4.67 (5.36) | 5.33 (5.24) |
| Age (years) | 60.94 (15.66) | 61.04 (15.77) | 60.38 (15.9) | 60.67 (14.47) |
| Baseline MD (dB) | -3.55 (5.71) | -3.57 (5.72) | -3.76 (6.12) | -3.23 (5.17) |
| Baseline PSD (dB) | 3.42 (3.29) | 3.42 (3.27) | 3.49 (3.41) | 3.42 (3.34) |

MD = mean deviation; PSD = Pattern standard deviation. Values are presented as mean (standard deviation), unless otherwise noted.

### A Deep Variational Autoencoder for Visual Fields

We utilized a VAE to create a generative process for visual field data. A VAE is an unsupervised learning method that transforms high-dimensional data to a small set of latent features^17^. Furthermore, the advantage of the VAE is that the latent features can be disentangled to generate synthetic visual fields^20^. The VAE was derived from variational Bayesian methods^21^, and the model is optimized to have minimal image reconstruction loss, while maintaining a homogenous feature space based on a regularization prior. In vanilla VAE, for regularization the Kullback-Leibler divergence^22^ is used, however for visual field data we use the maximum mean discrepancy^23^.

VAEs consist of two components, the encoder and decoder. The encoder is a function that takes in a visual field and compresses it to a lower-dimensional space. The decoder (sometimes called the generator) is a probabilistic distribution that maps the latent features back to the original high-dimensional image space. Since the decoder is trained to reconstruct the original data using a small number of latent dimensions, those features end up representing key aspects of the data. In other words, the encoder-decoder process is forced to learn the most important features of the original data. Furthermore, because there is a regularization prior, the VAE learns features that are homogenous, and therefore can be interpreted clinically. The bottleneck structure of the VAE is presented in Figure 1A.

Both the encoder and decoder used neural networks with deep learning. The encoder is itself a network, but for the decoder, the network is the mean of a Gaussian distribution. Details of both networks can be found in Figure 1B. The encoder of the VAE is comprised of two 2D convolutional layers, a reshaping layer and a fully-connected dense layer. The decoder begins with a fully-connected dense layer, followed by a reshaping layer, and then a sequence of deconvolutions, which up-sample until the original 12 × 12 dimension is reached. All layers use a 3 × 3 kernel size and a stride of two, and the activation is a rectified non-linear unit transformation, except for F1 and D3, which used the identity and sigmoid activations, respectively.

For each choice of latent dimension, the model was trained using the Adam optimizer, an extension of stochastic gradient descent, using 50 epochs and a batch size of 100^24^; we used a learning rate of 1e-4. The training epoch with the minimal validation loss was chosen as optimal. The VAE was implemented using the deep learning library Keras (version 2.2)^25^ with Tensorflow (version 1.9)^26^ backend, all within RStudio (3.5.1)^27^.

### Dimension of the Latent Features

The dimension of the latent space is a parameter that was specified to reflect the number of latent features that can adequately explain the high-dimensional visual fields. In this study, we explored the performance of the VAE over latent dimensions ranging from one to ten. Throughout the analysis all results based on the VAE are reported for all ten variations of the latent dimension in an attempt to determine the preferred latent space.

### Rates of Visual Field Progression

We used the latent features of the VAE to determine rates of progression for glaucoma patients in the test dataset. For each patient, we obtained the latent features corresponding to each visual field using the encoder. Then, we studied the longitudinal trends of the latent features (instead of the complex 52-dimensional visual field). In particular, we quantified progression based on the global rate of change across all features from a model. In order to calculate this global rate, we performed ordinary least squares (OLS) regression with a zero-sum constraint on the design matrix; thus, allowing us to perform a hypothesis test on the mean rate of change across time. Progression was defined as a significant global rate of change based on a two-sided hypothesis test with type-1 error of 0.05.

This rate of visual field progression can be interpreted as the speed of deterioration for a patient’s visual field features, where features represent some lower dimensional representation of the visual field. This is reminiscent of SAP MD, which is a comparison of a current visual field to an age adjusted healthy baseline in one-dimension. As such, for comparison we used standard rates of visual field progression of MD using OLS linear regression across time, with progression defined as a significantly negative rate of change over time (alpha = 0.05).

Finally, in the absence of a gold standard of progression, we compared the progression methods by matching their specificities at 95%. Because both methods are p-values with a type-1 error of 0.05, this is automatically achieved with a cutoff of 0.05. Therefore, the method with the higher rate of detecting progression in the glaucoma patients in the test dataset has superior operating characteristics. Note that this is not a sensitivity, because not all patients are progressing, but rather a progression hit rate. For each method, we estimated hit rate percentage at two, four and six years from baseline visits and also for all patient follow-up. Only visits that occurred on or before the cutoff time are included in the analysis. This imitated a clinical setting where each metric is calculated at every visit and progression is diagnosed. For hypothesis testing, bootstrapped confidence intervals were presented^28^.

### Predicting Future Visual Fields

An advantage to the VAE modeling framework is the ability to generate visual fields from its latent features through the decoder. This motivated a method for predicting future visual fields through a two-stage procedure. First, for a longitudinal collection of visual fields, we obtained their corresponding latent features (obtained from the encoder) and modeled each dimension independently using OLS linear regression. Then, we used the predicted values of the latent features to generate future visual fields using the decoder. Note that because the decoder is a mean process, the generated visual fields are de-noised.

We assessed the prediction accuracy of this two-stage approach by predicting the fourth, sixth, and eighth follow-up visits from the first three visits of all patients in the test dataset. We also looked at prediction accuracy in glaucoma patients only. Prediction accuracy was assessed using mean absolute error (MAE) for only the 52 informative locations (i.e., not the full 12 × 12 image). All results are compared to the established technique of visual field prediction, PW linear regression. One-sided Wilcoxon signed rank tests are presented to formally compare if the MAE from the VAE is smaller than from PW, using a Bonferroni corrected type-1 error, 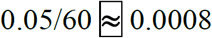.

## RESULTS

This study included 29,701 visual fields; however, 540 visual fields were excluded due to false-positives greater than 15% or fixation error greater than 33%. This yielded 29,161 usable visual fields from 3,832 eyes. These patient eyes had an average follow-up of 4.95 years with a mean of 7.61 visits. Mean age at baseline was 60.94, mean baseline MD was −3.55 dB and mean baseline PSD was 3.42. From the usable visual fields, we created training, validation, and test datasets, by randomly sampling eyes with probability 80%, 10%, and 10%, respectively. Population characteristics for each dataset are presented in Table 1.

The models were trained for 50 epochs and the optimal losses are presented in Figure 2 for the training and validation datasets across models with varying latent dimensions. From the figure it is clear that the reconstruction losses were nearly zero for all models and that the regularization loss decreased with the number of dimensions after three dimensions.

**Figure 2.**
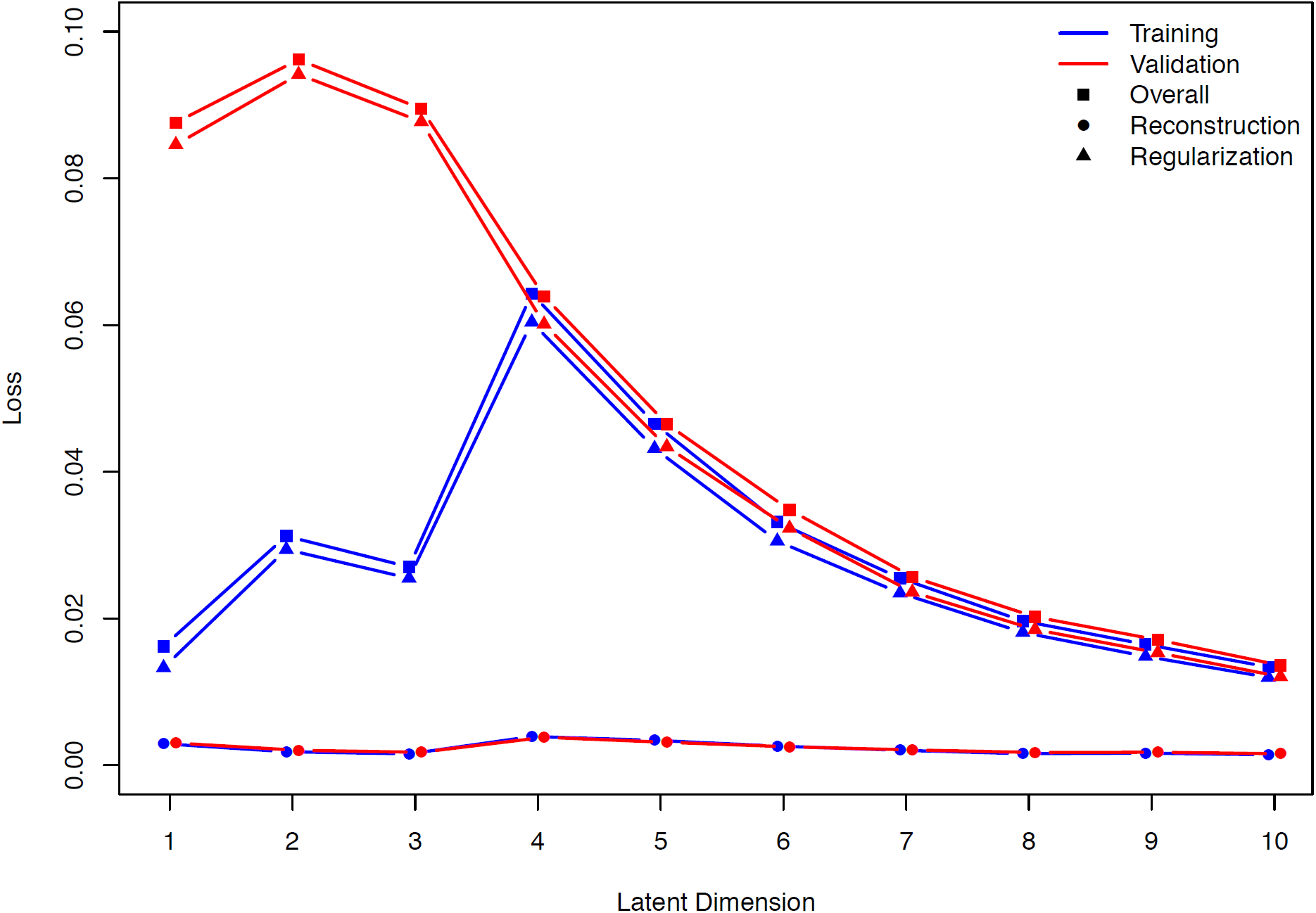
Training and validation loss over different latent dimensions. Overall loss is presented along with reconstruction and regularization loss.

We investigated the rates of visual progression determined from the latent space of the VAE and SAP MD. To motivate using the VAE for detecting progression, we have presented the longitudinal follow-up for examples patients of each disease status in a two-dimensional latent space (Figure 3A). The predicted trajectories are presented as arrows and the observed longitudinal follow-up are underlaid using muted colors. Furthermore, the location of a healthy visual field (blue square) is displayed, where healthy is defined as the mean visual field of all the healthy eyes. In Figure 3B-C, the first and second latent feature are shown across time with OLS regression lines indicating rates of change for each feature. Our defined VAE progression metric is the average of feature slopes, which is a calculation of the speed of movement through the latent space.

**Figure 3.**
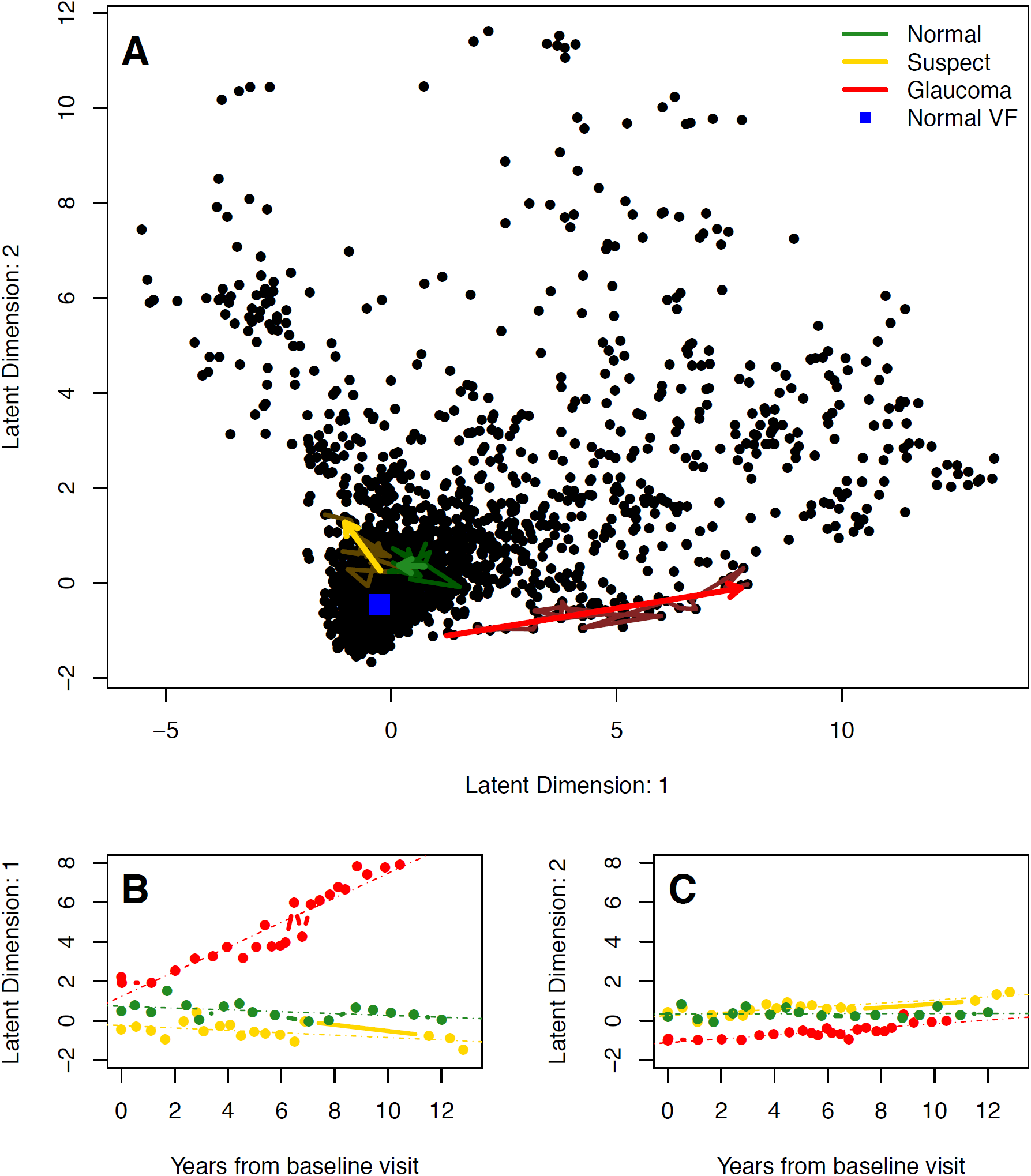
Visualizing the latent space of the VAE in two dimensions. A. The top frame shows predicted trajectories through the latent space, presented as arrows, are presented for example patients of each disease status. Underlaying the trajectories are the observed longitudinal follow-up for each patient using muted colors. The healthy visual field in latent space is a blue square near the origin. B. The bottom left frame shows the longitudinal latent features from the first dimension for the example patients. OLS regression lines are presented, which indicate rates of change in each latent dimension. C. The second dimension is shown in the bottom right.

In Figure 4, the progression detection hit rates are presented for of each of the diagnostic measures. The vertical line in each frame is a representation of the 95^th^ percentile of MD, and thus metrics with no overlap are significantly superior. At two years from baseline, the VAE models with five, seven, and eight dimensions were significantly superior at detecting progression; while at four years the VAE models with seven and eight dimensions remained significant. The significance disappears by six years, however the VAE models with seven and eight dimensions remain superior.

**Figure 4.**
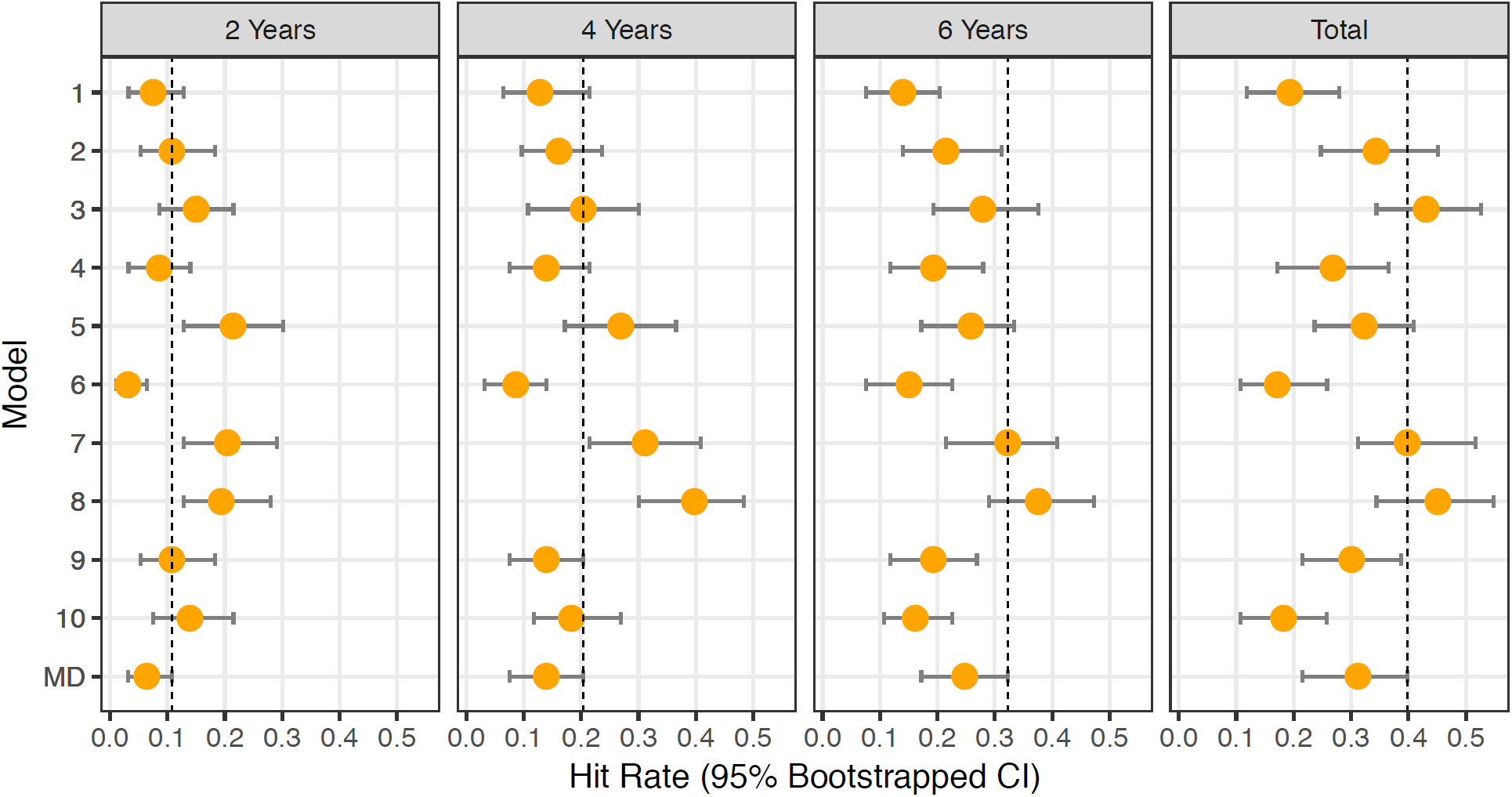
Visual field progression hit rate for diagnostic measures at two, four, and six years from baseline visit, and total follow-up, for glaucoma patients in the test dataset. Error bands represent 95% bootstrapped confidence intervals.

To verify that the trained models effectively learned the generative process we assessed the predictive capacity and the results are presented in Figures 5 and 6, plus Table 2. Figure 5 shows boxplots of the MAE for all patients in the test dataset along with glaucoma patients only. The MAE were most similar between VAE and PW when predicting the fourth visit; however, the VAE is generally superior with only the model with one dimension not significantly superior (P = 0.1528), with the rest having P-values less than 0.0001. A similar pattern holds for glaucoma patients, however, now only the VAE models with more than three latent dimensions were significantly superior to PW. Superiority of the VAE prediction increased when predicting further into the future, to visits six and eight, as all VAE models had significantly superior prediction, for all patients and those with glaucoma (Table 2).

**Table 2.** Mean absolute error (MAE) (in dB) over latent dimensions, along with the mean across dimensions and point-wise (PW) linear regression. Predictions are of the fourth, sixth, and eighth follow-up visits, using the first three visits from all patients in the test dataset and patients with glaucoma at baseline. Summaries of MAE are presented only for the 52 informative locations (i.e., not the full 12×12 image). P-values correspond to the one-sided Wilcoxon signed rank test, comparing each latent dimension to the PW prediction. The Bonferroni corrected type 1 error is 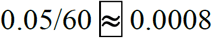.

|  | Model | Visit 4 |  | Visit 6 |  | Visit 8 |  |
| --- | --- | --- | --- | --- | --- | --- | --- |
|  |  | Median (95% CI) | P-value | Median (95% CI) | P-value | Median (95% CI) | P-value |
| All Patients | 1 | 2.83 (1.09, 12.10) | 0.1528 | 3.29 (1.09, 14.32) | <0.0001 | 3.76 (1.11, 13.93) | <0.0001 |
|  | 2 | 2.45 (1.07, 10.94) | <0.0001 | 3.08 (1.12, 12.52) | <0.0001 | 3.54 (1.22, 17.50) | <0.0001 |
|  | 3 | 2.43 (1.08, 11.14) | <0.0001 | 2.92 (1.15, 13.31) | <0.0001 | 3.48 (1.16, 20.05) | <0.0001 |
|  | 4 | 2.52 (1.07, 12.41) | <0.0001 | 3.05 (1.23, 14.35) | <0.0001 | 3.60 (1.24, 17.79) | <0.0001 |
|  | 5 | 2.42 (1.03, 13.39) | <0.0001 | 3.15 (1.13, 15.29) | <0.0001 | 3.37 (1.15, 18.16) | <0.0001 |
|  | 6 | 2.38 (1.01, 11.23) | <0.0001 | 3.18 (1.15, 16.25) | <0.0001 | 3.81 (1.22, 21.83) | <0.0001 |
|  | 7 | 2.46 (1.08, 10.20) | <0.0001 | 3.32 (1.21, 17.50) | <0.0001 | 4.04 (1.34, 20.01) | <0.0001 |
|  | 8 | 2.50 (1.09, 10.56) | <0.0001 | 3.31 (1.26, 18.55) | <0.0001 | 4.06 (1.31, 22.74) | <0.0001 |
|  | 9 | 2.51 (1.02, 12.76) | <0.0001 | 3.26 (1.29, 15.59) | <0.0001 | 3.81 (1.37, 21.06) | <0.0001 |
|  | 10 | 2.83 (1.12, 12.06) | <0.0001 | 3.58 (1.29, 19.37) | <0.0001 | 4.42 (1.43, 25.98) | <0.0001 |
|  | Mean | 2.53 (1.07, 11.68) | -- | 3.21 (1.19, 15.70) | -- | 3.79 (1.25, 19.90) | -- |
|  | PW | 3.12 (1.40, 21.50) | -- | 4.27 (1.86, 29.65) | -- | 6.06 (2.33, 65.08) | -- |
| Glaucoma Only | 1 | 5.11 (1.58, 16.13) | 0.8411 | 5.46 (1.91, 19.00) | <0.0001 | 5.52 (1.76, 14.69) | <0.0001 |
|  | 2 | 4.84 (1.73, 10.94) | 0.5878 | 5.04 (2.03, 13.35) | <0.0001 | 6.04 (2.08, 18.98) | <0.0001 |
|  | 3 | 4.37 (1.51, 10.89) | 0.0832 | 4.93 (2.06, 15.04) | <0.0001 | 5.50 (2.08, 21.17) | <0.0001 |
|  | 4 | 4.18 (1.55, 12.41) | 0.0018 | 4.86 (2.17, 16.15) | <0.0001 | 5.59 (1.79, 18.80) | <0.0001 |
|  | 5 | 4.07 (1.67, 12.34) | 0.0002 | 4.96 (1.96, 15.99) | <0.0001 | 5.27 (2.06, 20.29) | <0.0001 |
|  | 6 | 4.02 (1.52, 10.79) | <0.0001 | 5.13 (1.80, 18.60) | <0.0001 | 5.51 (1.86, 23.65) | <0.0001 |
|  | 7 | 4.14 (1.60, 9.98) | 0.0001 | 5.44 (1.82, 18.09) | <0.0001 | 5.92 (1.74, 20.52) | <0.0001 |
|  | 8 | 3.87 (1.57, 10.26) | <0.0001 | 4.62 (1.79, 19.43) | <0.0001 | 5.93 (2.05, 25.58) | <0.0001 |
|  | 9 | 4.04 (1.54, 12.28) | <0.0001 | 4.92 (1.83, 15.81) | <0.0001 | 5.84 (1.61, 22.61) | <0.0001 |
|  | 10 | 4.33 (1.52, 12.06) | 0.0001 | 4.94 (2.04, 20.99) | <0.0001 | 5.54 (2.27, 27.59) | <0.0001 |
|  | Mean | 4.30 (1.58, 11.81) | -- | 5.03 (1.94, 17.24) | -- | 5.67 (1.93, 21.39) | -- |
|  | PW | 4.24 (2.01, 21.50) | -- | 6.69 (2.85, 52.96) | -- | 7.66 (2.90, 85.13) | -- |

**Figure 5.**
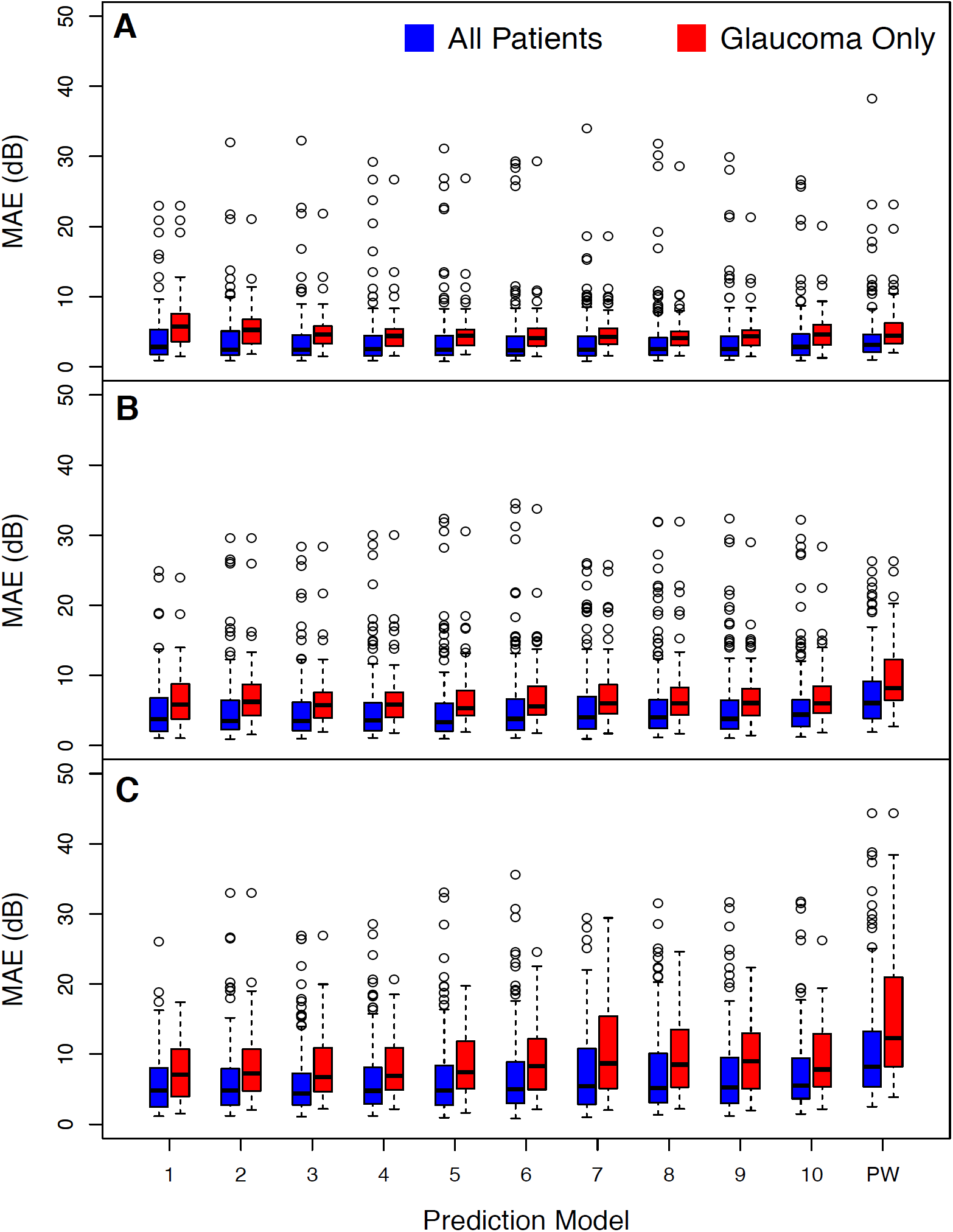
Comparing the prediction accuracy of the VAE and PW methods. Boxplots present summaries of prediction using MAE (dB) over the varying number of latent dimensions. A. Predictions in the top frame are of the fourth follow-up visit, using the first three visits from all patients in the test dataset (blue) and glaucoma patients only (red). Summaries of MAE are presented only for the 52 informative locations (i.e., not the full 12×12 image). Due to the upper-bound of the y-axis, there are 17 outliers that have been truncated from the PW model across the three frames. B. Predictions for the sixth visit are in the middle frame. C. Predictions for the eighth visit are in the bottom frame.

**Figure 6.**
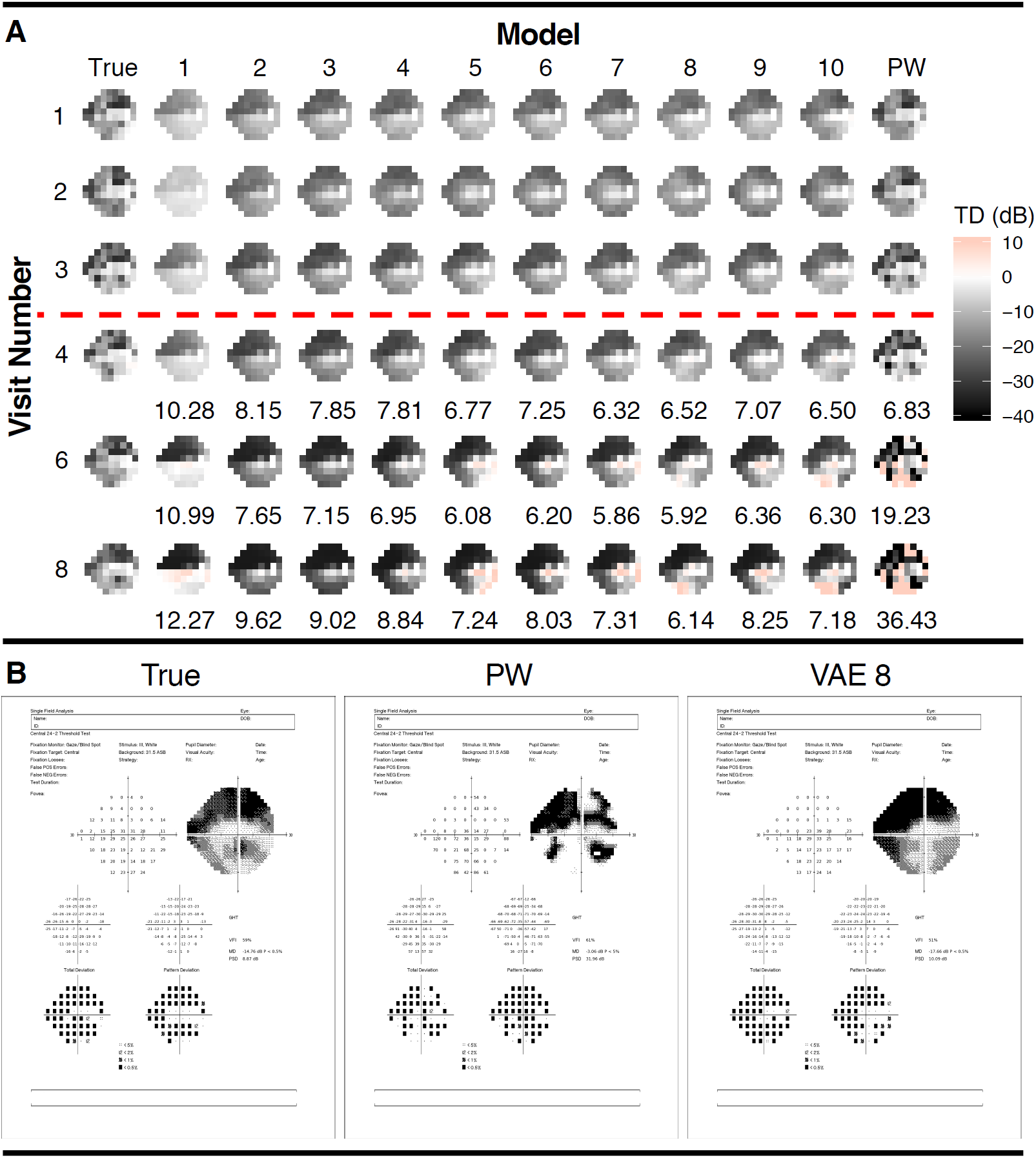
Demonstrating prediction capability of the VAE compared to the PW method using a glaucoma patient in the test dataset. A. In the top frame accompanying the predicted visual fields are MAE (dB) from the fourth, sixth, and eighth follow-up visits, using the first three visits from baseline. The first three visual fields represent fitted output from the fitted models. B. The bottom frame presents the predictions at the eighth visit using the Zeiss printout. The VAE model presented is with eight latent dimensions.

The ability of the VAE to predict future visual fields is further illustrated in Figure 6, where predictions are presented of an example glaucoma patient with severe disease. The predictions from the VAE model are stable when predicting future patterns of vision loss, while PW deteriorates, with a MAE of 36.43 when predicting the eighth visit. In Figure 6B, the predicted visual field at the eighth visit is presented using the Zeiss printout, for the true visual field, PW and the VAE model with eight dimensions.

To further demonstrate the clinical utility of the VAE, we visualized it with a two-dimensional latent space across clinical measures. Figure 7 shows the relationship between the VAE latent feature space and clinical variables, disease status, age, MD, and PSD. Finally, in Figure 8, we presented the generative distribution derived from the VAE for two latent dimensions, with the predicted trajectories of each of the example patients included. Overlaying each path is a blue line that indicates the distance traveled in the first year of follow-up, which is constant across time.

**Figure 7.**
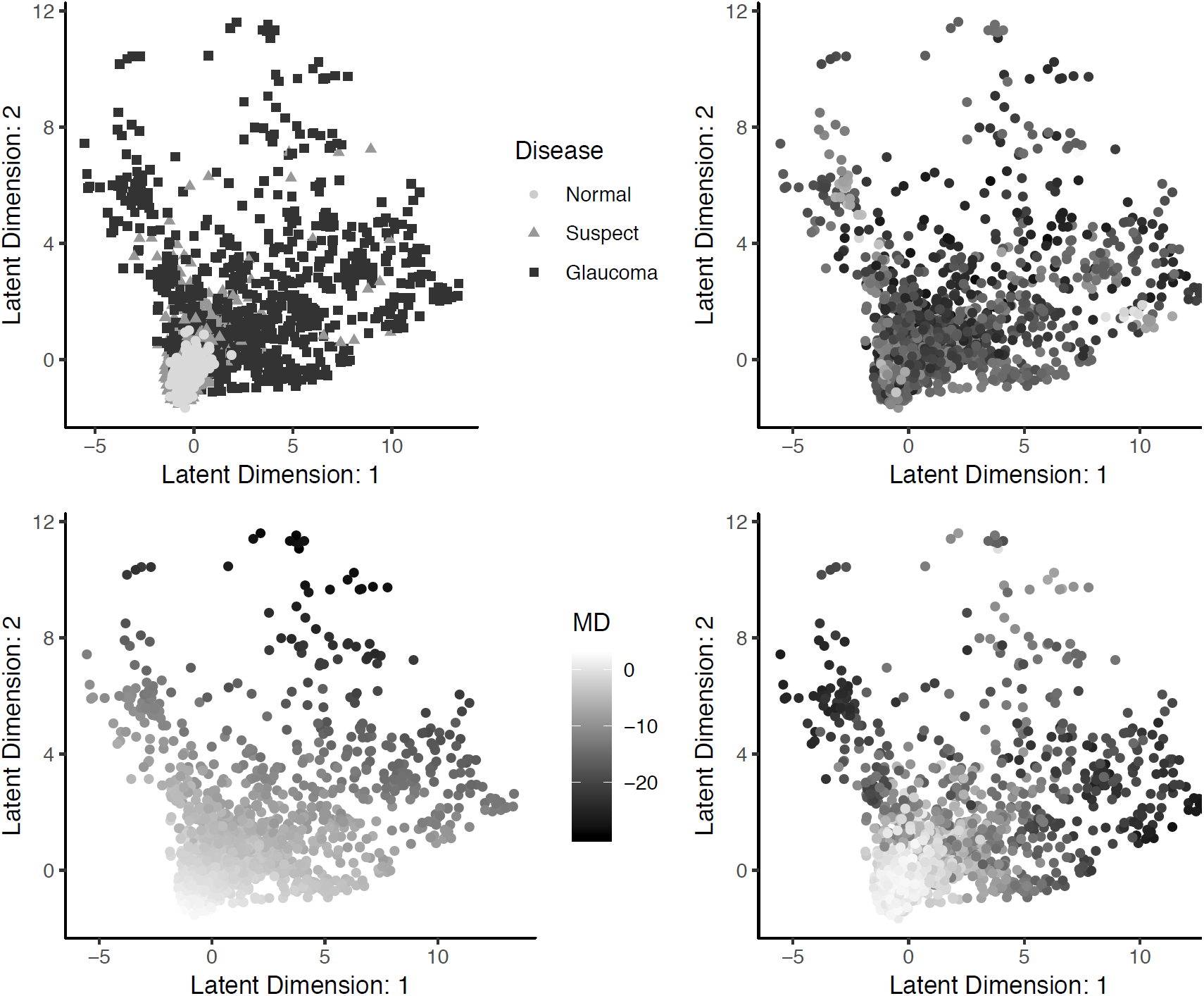
Visualization of the relationship between the VAE latent feature space in two dimensions and clinical variables, disease status, age, MD, and PSD.

**Figure 8.**
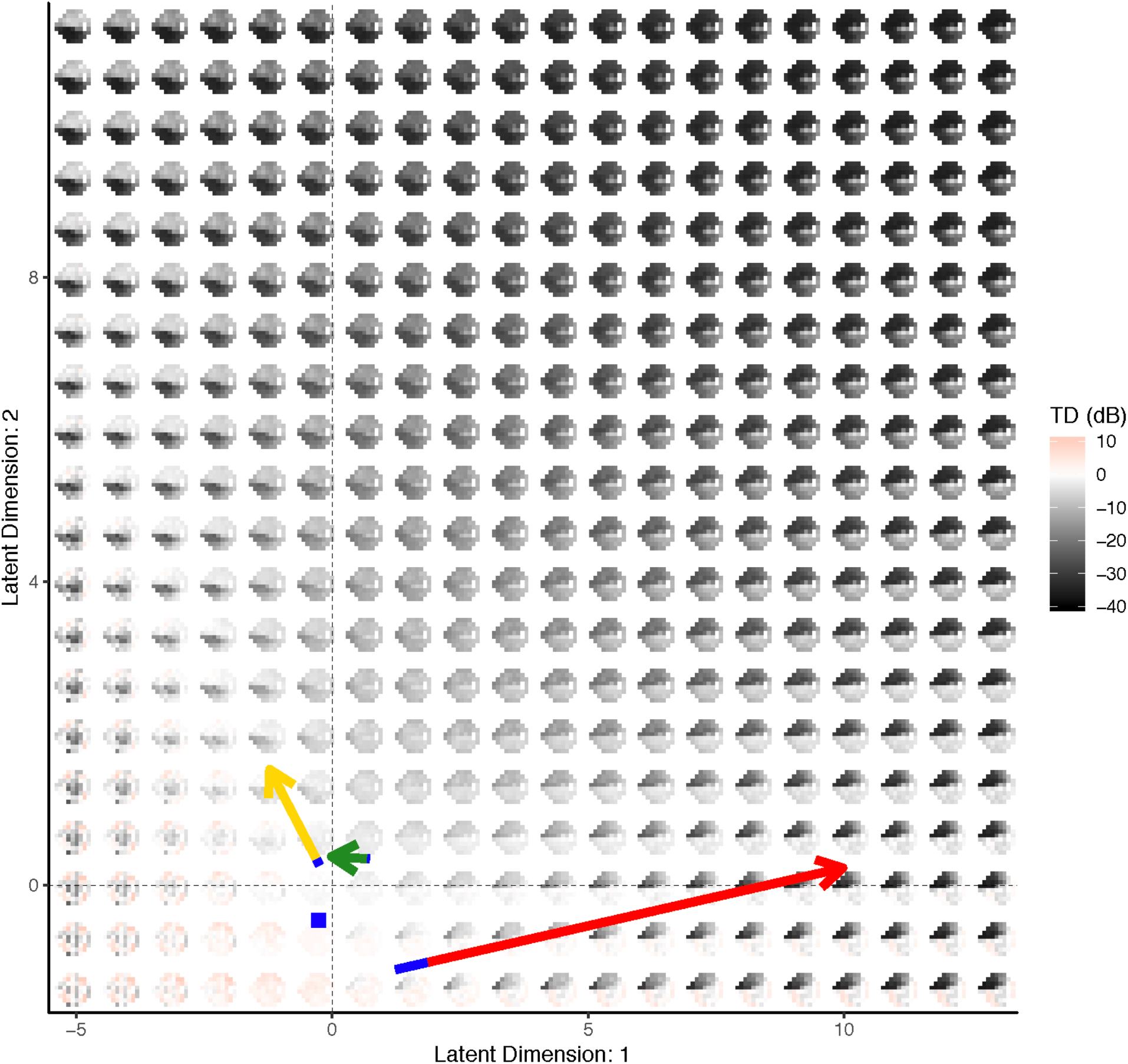
Visualizing the VAE generative distribution for visual fields with two latent dimensions. Included are the paths of the example patients from Figure 4. The length of the blue line overlaying each path indicates the speed of change across time and is the length traveled in the first year. The average visual field from all of the normal patients is represented as a blue square.

## DISCUSSION

In this manuscript, we developed and validated a novel deep learning algorithm that produced a unified framework for learning the generative process of visual fields and detecting rates of glaucoma progression. The VAE used a low-dimensional latent space representation of the more complex and high-dimensional visual field image to produce a clinically relevant latent space. This was achieved through the model structure of the VAE, which learned the latent features through both a compression (i.e., encoder) and generative (i.e., decoder) process. Standard techniques for learning latent features, such as probabilistic principle components, factor analysis or independent components analysis, only include the compression component and, consequently, the latent space can often be less realistic^7, 29^. Furthermore, these standard techniques assume the latent features are a linear combination of the original image, while the VAE uses deep learning to allow for arbitrarily non-linear mappings.

Previous studies have used machine learning approaches to assess glaucomatous damage from visual fields^13, 14, 30, 31^. In these studies, the glaucomatous status of the eyes was determined either by clinical measures or by clinical expertise, making the machine learning performance dependent on the chosen gold standard. This type of technique can be limited, because the end-point of interest is often inaccurate or different from the true clinical end-point. We circumvented this issue in our study by using an unsupervised method. Instead of training against a noisy gold standard, the VAE trains a visual field against itself, in the process developing a generative model through clinically interesting latent features.

To visualize the latent features for visual fields, we presented the latent space in two-dimensions for clinical measures, glaucoma status, age, MD, and PSD (Figure 7). Through inspection of these figures the latent space comes to life. Across all clinical measures it is clear that patients with no evidence of glaucoma reside near the origin, as this is the location of the normal disease patients, and young patients with MD and PSD values near zero. As the latent features increase in either dimension the disease severity increases. In particular, both age and MD values appear to become more severe (i.e., older age and more negative MD) with an increase in either of the features or a combination of both. In contrast, PSD only demonstrates more severity (i.e., larger values) with an increase in either of the latent features, but not both. This illuminates a clinical phenomenon, that MD and age are linear risk factors, while PSD identifies local defects, which are typically not evident in the late stages of disease.

The latent space can also be used to visualize the generative process for visual fields. In Figure 8, the distribution of visual fields across the two-dimensional latent space was displayed. This presentation provides more clinical context to the latent space, as the generative process is a smooth representation of the variability in visual fields. In particular, we can see the patterns of disease severity, as an increase in the second latent feature appears to indicate a global worsening in the functional vision, while the first feature dictates worsening patterns in the superior (or inferior) hemisphere when moving to the right (or left) of the origin. Because the VAE produced a latent feature space that was clinically informative, it had utility for determining rates of progression.

In particular, we leveraged the clinical interpretation of the latent space to calculate progression rates by measuring the rate of a patient’s movement through the generative distribution. We formally defined the progression rate based on the average of the rates across features. This process was exemplified through the visual fields of a normal, suspect, and glaucoma patient in Figure 3. The movement of each of the patients through latent space was displayed, along with the trajectories of the features across time. The glaucoma patient has the highest rate, indicated by the large positive slope in the first feature. In Figure 8, the trajectory of each of the example patients overlays the generative process, with an indicator of the rate of movement through latent space (blue line). This novel presentation of the trajectories through the distribution of visual fields provided a literal clinical road map for making treatment decisions.

We provided evidence that the rate of change through latent space is superior to MD (Figure 4) at two and four years from baseline follow-up for a number of VAE models, in particular with eight dimensions. This is not surprising, as MD can be thought of as a one-dimensional latent space representation of a visual field^32^. Therefore, it follows that the performance of the VAE would improve upon MD. Although the VAE models do not significantly outperform MD in the long run, this finding is nonetheless important because detection of change and intervention in the early years of follow-up can reduce a patient’s likelihood of vision loss^33^.

Furthermore, the VAE can be used to predict the future location of a patient’s latent features (Figures 5 and 6, Table 2). This allows for accurate predictions of future visual fields, which can help illuminate patterns and severity of progression. Such predictions of whole visual fields are obviously not possible with a single global metric such as MD. Even if one uses PW regression, the VAE was shown to have superior prediction capabilities. The predictions in Figure 6 demonstrated the benefits of using the VAE as compared to PW regression. In particular, while the PW method was highly susceptible to local variability, the VAE produced stable (i.e., smooth) predictions. This resulted in predictions that were robust to the variability of individual entries and consequently, the true disease pattern visible is better illuminated.

The prediction capability of the VAE model, including patients with patterns in their visual fields, is reassuring, as global methods, like MD, typically fail at detecting localized defects^33^. The flexibility of the VAE, to not only improve rates of detecting progression over the global MD method, but also maintain superior predictions over PW regression, makes it clinically useful. In particular, because the VAE has the ability to detect progression at higher rates in the early years from baseline, clinicians will need fewer visits to obtain accurate detection of progression; thus, limiting the burden on patients.

A potential limitation, or motivation for future work, is that when training the deep learning model, we treated each of the visual fields as independent images. We overcame this limitation by modeling a patient’s longitudinal visual fields series in latent space, which we showed to be an effective method for assessing rates of progression. In the future, we could make improvements by learning a generative model for longitudinal series of visual fields, instead of a singular image. This would be a powerful technique for clustering patients, not by their visual fields, but by the characteristics of their progression.

Another extension to the method presented in this manuscript is to generate synthetic glaucomatous visual fields to be used as a benchmark dataset to validate new methods for glaucoma research. Existing methods attempt to generate longitudinal visual fields by modelling PW relationships across time with potentially non-linear fits^34, 35^, however they typically ignore spatial correlations in the visual field or can only predict stable fields. The decoder from the VAE can be used to simulate glaucomatous visual field series that accounts for spatial dependencies and the highly-nonlinear PW trends.

In conclusion, this manuscript showed the potential use of the VAE latent space for assessing rates and trajectories of glaucoma progression. The rates of progression can be considered a multi-dimensional extension of MD with improved abilities to detect progression and the additional benefit of a generative technique to predict future patterns and severity of visual fields.

## ACKNOWLEDGMENTS/DISCLOSURE

a. Funding/Support: This work was supported in part by National Institutes of Health/National Eye Institute [grant numbers EY029885 (FAM), EY027651 (FAM), and EY021818 (FAM)].
b. Financial Disclosures: **Samuel I. Berchuck**: none; **Sayan Mukherjee**: none; **Felipe A. Medeiros**: Alcon Laboratories (C, F, R), Allergan (C, F), Bausch&Lomb (F), Carl Zeiss Meditec (C, F, R), Heidelberg Engineering (F), Merck (F), nGoggle Inc. (F), Sensimed (C), Topcon (C), Reichert (C, R).
c. Other Acknowledgments: none.

## REFERENCES

1. Weinreb RN, Aung T, Medeiros FA. The pathophysiology and treatment of glaucoma: a review. JAMA 2014;311:1901–11.

2. Vianna JR, Chauhan BC. How to detect progression in glaucoma. Prog Brain Res 2015;221:135–58.

3. Chauhan BC, Garway-Heath DF, Goni FJ, et al. Practical recommendations for measuring rates of visual field change in glaucoma. Br J Ophthalmol 2008;92:569–73.

4. Gardiner SK, Mansberger SL, Demirel S. Detection of Functional Change Using Cluster Trend Analysis in Glaucoma. Invest Ophthalmol Vis Sci 2017;58:BIO180–BIO190.

5. Goldbaum MH, Lee I, Jang G, et al. Progression of patterns (POP): a machine classifier algorithm to identify glaucoma progression in visual fields. Invest Ophthalmol Vis Sci 2012;53:6557–67.

6. Goldbaum MH, Sample PA, Zhang Z, et al. Using unsupervised learning with independent component analysis to identify patterns of glaucomatous visual field defects. Invest Ophthalmol Vis Sci 2005;46:3676–83.

7. Sample PA, Boden C, Zhang Z, et al. Unsupervised machine learning with independent component analysis to identify areas of progression in glaucomatous visual fields. Invest Ophthalmol Vis Sci 2005;46:3684–92.

8. Yousefi S, Goldbaum MH, Balasubramanian M, et al. Learning from data: recognizing glaucomatous defect patterns and detecting progression from visual field measurements. IEEE Trans Biomed Eng 2014;61:2112–24.

9. Abe RY, Diniz-Filho A, Costa VP, Gracitelli CP, Baig S, Medeiros FA. The Impact of Location of Progressive Visual Field Loss on Longitudinal Changes in Quality of Life of Patients with Glaucoma. Ophthalmology 2016;123:552–7.

10. Fitzke FW, Hitchings RA, Poinoosawmy D, McNaught AI, Crabb DP. Analysis of visual field progression in glaucoma. British Journal of Ophthalmology 1996;80:40–48.

11. Diaz-Pinto A, Colomer A, Naranjo V, Morales S, Xu Y, Frangi AF. Retinal Image Synthesis and Semi-supervised Learning for Glaucoma Assessment. IEEE Trans Med Imaging 2019.

12. Medeiros FA, Jammal AA, Thompson AC. From Machine to Machine: An OCT-Trained Deep Learning Algorithm for Objective Quantification of Glaucomatous Damage in Fundus Photographs. Ophthalmology 2018.

13. Li Z, He Y, Keel S, Meng W, Chang RT, He M. Efficacy of a Deep Learning System for Detecting Glaucomatous Optic Neuropathy Based on Color Fundus Photographs. Ophthalmology 2018;125:1199–1206.

14. Yousefi S, Kiwaki T, Zheng Y, et al. Detection of Longitudinal Visual Field Progression in Glaucoma Using Machine Learning. Am J Ophthalmol 2018;193:71–79.

15. Wen JC, Lee CS, Keane PA, et al. Forecasting Future Humphrey Visual Fields Using Deep Learning. arXiv preprint arXiv:180404543 2018.

16. Goodfellow I, Pouget-Abadie J, Mirza M, et al. Generative adversarial nets. In Advances in neural information processing systems 2014.

17. Kingma DP, and Max Welling. Auto-encoding variational Bayes. arXiv preprint arXiv:13126114 2013.

18. Way GP, Greene CS. Extracting a biologically relevant latent space from cancer transcriptomes with variational autoencoders. Pacific Symposium on Biocomputing Pacific Symposium on Biocomputing 2018;23:80–91.

19. Choi H, Kang H, Lee DS, Alzheimer’s Disease Neuroimaging I. Predicting Aging of Brain Metabolic Topography Using Variational Autoencoder. Front Aging Neurosci 2018;10:212.

20. Doersch C. Tutorial on variational autoencoders. arXiv preprint arXiv:160605908 2016.

21. Blei DM, Kucukelbir A, McAuliffe JD. Variational Inference: A Review for Statisticians. Journal of the American Statistical Association 2017;112:859–877.

22. Kullback S, Leibler RA. On Information and Sufficiency. The Annals of Mathematical Statistics 1951;22:79–86.

23. Zhao S, Jiaming Song, and Stefano Ermon. InfoVAE: Information maximizing variational autoencoders. arXiv preprint arXiv:170602262 2017.

24. Kingma DP, Ba J. Adam: A method for stochastic optimization. arXiv preprint arXiv:14126980 2014.

25. Chollet F, others. Keras: GitHub, 2015.

26. Abadi M, Agarwal A, Barham P, et al. TensorFlow: Large-scale machine learning on heterogeneous systems. Software available from tensorfloworg 2015.

27. Chollet F, Allaire J, others. R Interface to Keras. https://github.com/rstudio/keras: GitHub, 2017.

28. Efron B, Tibshirani RJ. An Introduction to the Bootstrap: CRC press, 1994.

29. Wetzel SJ. Unsupervised learning of phase transitions: From principal component analysis to variational autoencoders. Phys Rev E 2017;96:022140.

30. Asaoka R, Murata H, Iwase A, Araie M. Detecting Preperimetric Glaucoma with Standard Automated Perimetry Using a Deep Learning Classifier. Ophthalmology 2016;123:1974–80.

31. Kim SJ, Cho KJ, Oh S. Development of machine learning models for diagnosis of glaucoma. PLoS One 2017;12:e0177726.

32. Leske MC, Heijl A, Hyman L, et al. Predictors of long-term progression in the early manifest glaucoma trial. Ophthalmology 2007;114:1965–72.

33. Bengtsson B, Heijl A. A visual field index for calculation of glaucoma rate of progression. Am J Ophthalmol 2008;145:343–53.

34. Russell RA, Garway-Heath DF, Crabb DP. New insights into measurement variability in glaucomatous visual fields from computer modelling. PLoS One 2013;8:e83595.

35. Wu Z, Medeiros FA. Development of a Visual Field Simulation Model of Longitudinal Point-Wise Sensitivity Changes From a Clinical Glaucoma Cohort. Transl Vis Sci Technol 2018;7:22.

